# Dopamine modulates learning-related changes in dynamic striatal-cortical connectivity in Parkinson’s disease

**DOI:** 10.1101/619478

**Authors:** Raphael T. Gerraty, Madeleine E. Sharp, Amanda Buch, Danielle S. Bassett, Daphna Shohamy

## Abstract

Learning from reinforcement is thought to depend on striatal dopamine inputs, which serve to update the value of actions by modifying connections in widespread cortico-striatal circuits. While considerable research has described the activity of individual striatal and midbrain regions in reinforcement learning, the broader role for dopamine in modulating network-level processes has been difficult to decipher. To examine whether dopamine modulates circuit-level dynamic connectivity during learning, we characterized the effects of dopamine on learning-related dynamic functional connectivity estimated from fMRI data acquired in patients with Parkinson’s disease. Patients with Parkinson’s disease have severe dopamine depletion in the striatum and are treated with dopamine replacement drugs, providing an opportunity to compare learning and network dynamics when patients are in a low dopamine state (off drugs) versus a high dopamine state (on drugs). We assessed the relationship between dopamine and dynamic connectivity while patients performed a probabilistic reversal learning task. We found that reversal learning altered dynamic network flexibility in the striatum and that this effect was dependent on dopaminergic state. We also found that dopamine modulated changes in connectivity between the striatum and specific task-relevant visual areas of inferior temporal cortex, providing empirical support for theories stipulating that value is updated through changes in cortico-striatal circuits. These results suggest that dopamine exerts a widespread effect on neural circuitry and network dynamics during reinforcement learning.

## Introduction

Updating actions based on feedback is critical for survival in a changing environment. Accordingly, reinforcement has been central to our understanding of learning in psychology and neuroscience (Thorndike, 1898; Skinner, 1948; Schultz et al., 1997). Learning from reinforcement relies on the formation of associations between sensory cues, actions, and the value of outcomes, which must involve coordinated routing and processing of information across widespread brain areas. However, much of the research examining the neural and cognitive mechanisms of reinforcement learning has focused on describing the roles of individual brain regions. Despite the fact that circuit-level changes play a major role in theories of how value is updated in reinforcement learning (White, 1989b; Glimcher, 2011), there has been surprisingly little empirical work characterizing these circuit-level dynamics.

Studies of reinforcement learning have primarily focused on the role of the striatum and its input from midbrain dopamine neurons. The striatum has long been theorized to play an integrative role in brain function due to its widespread projections to cortical areas and its output to the motor system (Kemp and Powell, 1971; Bogacz and Gurney, 2007; Hikosaka et al., 2014; Ding, 2015). The role of the striatum in behavioral selection is thought to depend on its diverse inputs and its ability to gate and amplify outputs through a series of cortico-striatal loops (Yin and Knowlton, 2006).

It has been demonstrated repeatedly that the striatum is necessary for learning from reinforcement (White, 1989a; Robbins and Brown, 1990; Yin et al., 2006; Vo et al., 2014) and that intact striatal function depends on dopaminergic inputs from the midbrain (Kao and Powell, 1986; Westerink and Kwint, 1996; Steinberg et al., 2013). The precise learning signal carried by midbrain dopamine neurons has been well characterized at both the computational and physiological levels as a “reward prediction error” that signals the difference between received and expected rewards (Schultz et al., 1997; Sutton and Barto, 1998; Pagnoni et al., 2002; Bayer and Glimcher, 2005; Daw et al., 2006). It is theorized that this dopaminergic signal modulates cortico-striatal circuits (Glimcher, 2011). However, so far there has been little evidence in humans linking dopaminergic modulation with learning-related changes in striatal-cortical connectivity. More generally, while reinforcement learning theories in neuroscience posit a widespread circuit-level mechanism, studies of the time-varying interactions between distributed regions during learning have been lacking.

This lack of evidence linking dopamine, reinforcement learning, and circuit-level interactions has been due in part to a dearth of methods for testing the existence and nature of interregional interactions. While measures of the activation of individual regions provide a dynamic portrait of brain processes, most measures of interactions between regions have been based on pairwise structure in static correlations of regional activity time series. Recent studies in network neuroscience have begun to address these difficulties using emerging tools from graph theory to characterize the evolution of dynamic connectivity patterns over the same time scales as behavior change (Bassett et al., 2011; Bassett et al., 2015; Shine et al., 2016). More specifically, recent studies have begun to apply this approach to reinforcement learning, demonstrating that dynamic and large-scale connectivity centered on the striatum is associated with learning at the behavioral level and in model-derived estimates of learning (Mattar et al., 2016; Gerraty et al., 2018). However, these studies leave open two critical questions. First, while they show a correlation between the flexibility of dynamic network connectivity centered on the striatum and learning, they do not assess whether changes to the learned value of stimuli drive changes in network flexibility. And critically, these studies were not able to assess whether any observed changes in learning and network flexibility are related to dopamine.

A powerful approach to assessing dopaminergic modulation of learning comes from studies of patients with Parkinson’s disease. Patients with Parkinson’s suffer from a loss of striatal dopamine, which has been linked in some studies to impairments in reinforcement learning (Knowlton et al., 1996; Frank et al., 2004; Palminteri et al., 2009; Rutledge et al., 2009; Foerde et al., 2012; Schmidt et al., 2014; Sharp et al., 2015). The main treatment for Parkinson’s disease is levodopa, a dopamine precursor that leads to increased levels of dopamine in the striatum. Medication manipulations in patients with Parkinson’s disease have been used to model dopamine’s effects on learning by comparing behavioral performance and brain activity when patients are on versus off medication.

We used this approach in the current study, testing Parkinson’s patients on vs. off dopaminergic medication to examine dopamine’s effect on dynamic network connectivity in the striatum during reinforcement learning. Given prior findings demonstrating a relationship between dynamic cortico-striatal connectivity and value learning, and the success of reinforcement learning theory in describing dopaminergic inputs to the striatum, we hypothesized that dopamine would modulate learning-related changes in dynamic connectivity between the striatum and task-relevant cortical areas. Specifically, we predicted that (i) reinforcement learning would involve flexible network connectivity changes in cortico-striatal circuits, (ii) dopamine would modulate these learning-related connectivity changes, and (iii) dopamine would specifically alter dynamic connectivity between the striatum and cortical regions involved in processing task-relevant sensory information.

We tested these predictions in patients with Parkinson’s disease undergoing fMRI while engaged in a reinforcement learning task. The task used visual-category-specific choice options to allow us to track specific cortical sensory regions, and it involved a reversal, allowing us to decouple learning and flexibility from time-on-task. We compared behavior and dynamic connectivity metrics across sessions in which patients were tested on vs. off dopaminergic medication using a within-subject design. Our findings indicate that learning-induced changes in dynamic striatal-cortical connectivity are modulated by dopamine and that these changes are most pronounced in connections with task-specific sensory regions, providing a link between reinforcement learning and network mechanisms underlying the acquisition of learned behavior.

## Methods

### Patients

Participants were thirty patients with idiopathic Parkinson’s disease (7 females, mean + SD age: 61.86 + 6.25 years). Patients were recruited from the Center for Parkinson’s Disease and other Movement Disorders at Columbia University Medical Center and from the Michael J. Fox Foundation Trial Finder website. All patients provided informed consent and were compensated $100 per day for taking part in the study. All aspects of the study were approved by Columbia University’s Institutional Review Board.

Parkinson’s patients were in the mild-to-moderate stage of the disease, as rated on the Unified Parkinson’s Disease Rating Scale (UPDRS) while off medication by a neurologist specializing in movement disorders (mean + SD: 20.35 + 9.22). Two patients were not rated and five other patients were missing ratings for one session due to the lack of an available neurologist. The range of disease duration was 1-17 years. All patients had been receiving levodopa treatment for at least six months and the mean total daily levodopa or equivalent dose was 796.76 + 404.91 mg. Fifteen patients were also taking dopamine agonists. In addition to the learning task described below, patients also completed the Montreal Cognitive Assessment (MoCA), the forward and reverse digit span, Starkstein Apathy Scale, and a Beck Depression Inventory. Patients did not exhibit dementia (i.e. MoCA scores were > 26) and had no other history of major neurological or psychiatric illness except for Parkinson’s disease.

### Medication state

Each participant was tested in two sessions 24 hours apart and counterbalanced for order of medication state. For the OFF session, patients were asked to undergo an overnight withdrawal from all medication taken for Parkinson’s, lasting at least 16 hours in duration which is at least 10 half-lives for levodopa and 2 half-lives for dopamine agonists. In the ON session, the same patients were tested 1-1.5 hours after taking their normal dose of levodopa. To isolate the effect of levodopa, patients who were additionally taking dopamine agonists were asked to take only levodopa for the ON testing day. A comparison of UPDRS scores between OFF and ON sessions confirmed the expected effect of medication withdrawal on motor control (mean + standard error ON versus OFF difference: −10.48 + 0.96, t(22) = −10.87, *p* < 0.0001).

### Task

We designed a reinforcement learning task with two broad goals (**Figure 1**). First, we included a reversal of the optimal choice during each session, in order to manipulate learned choice contingencies and to characterize the effect of this manipulation on dynamic connectivity. Second, to characterize the dynamics of specific striatal-cortical circuits during learning, we used choice options that were images of objects and scenes, categories known to evoke specific patterns of activation in distinct regions of the ventral visual stream (Epstein and Kanwisher, 1998; Epstein et al., 1999; Reddy and Kanwisher, 2006). We predicted that the use of these categories during learning would lead to changes in striatal connectivity with domain-specific sensory areas in visual cortex.

**Figure 1.**
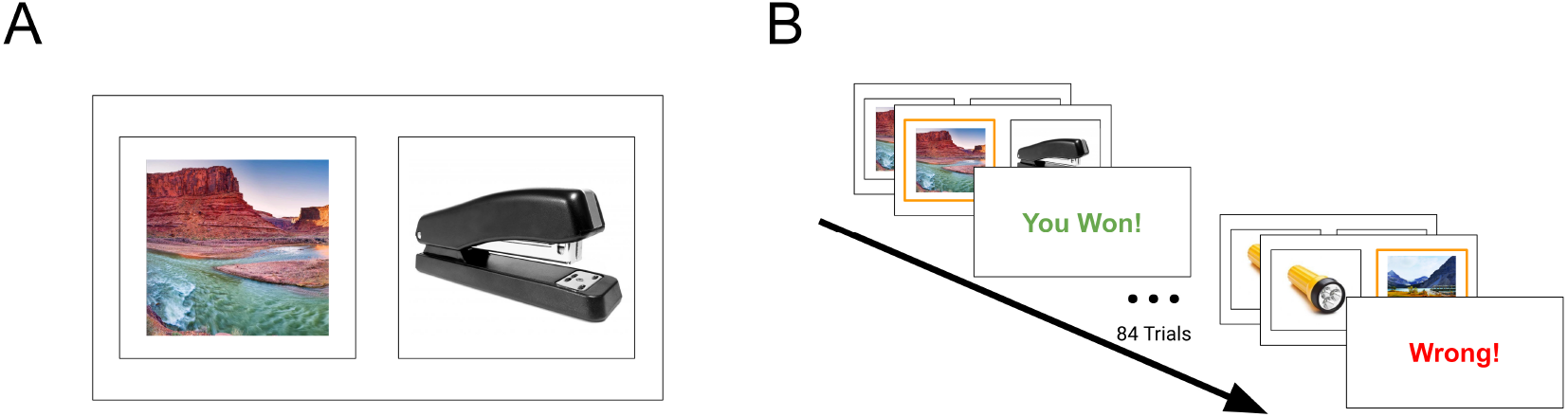
Learning task. **A)** On each trial, participants were asked to choose between an image of a scene and an image of an object. Each trial contained unique images that were randomly placed on the left or right side of the screen, and the participants indicated their decision with a button press. **B)** One category led to positive feedback with a probability of 0.7, while the other category led to positive feedback with a probability of 0.3. The category-outcome contingencies reversed after 85 trials.

On each trial, participants were asked to choose between images of a scene and of an object. The specific images varied on each trial and were randomly presented on the right and left sides of the screen. Participants responded using their index and middle fingers. There were 150 trials in each session (ON or OFF), broken up into five scan runs to allow short breaks for patients. Choosing the optimal category led to positive feedback (‘You Win!’) with a probability of 0.7, while choosing the non-optimal category led to positive feedback with a probability of 0.3. Negative feedback (‘Wrong!’) was shown with a probability of 0.3 and 0.7 for ‘correct’ and ‘incorrect’ choices respectively. Participants had 2.5 seconds to respond, followed by a 1.5-second crosshair and feedback shown for 1 second. A reversal took place on the 85th trial of each session, to allow participants enough time to learn the contingencies before the reversal. Participants were instructed that the correct option could change at any point during the task, and they were not told how many times a change could take place. All participants underwent a short practice prior to each scanning session in which simple shapes rather than image categories were used as stimuli, and experienced one reversal. Optimal categories at the start of the session were counterbalanced across subject-session pairs, such that half of the subjects experienced the same optimal category at the beginning of both sessions. Trials were separated by an intertrial interval (ITI) drawn from a truncated exponential distribution with a mean of 3 seconds and a minimum of 1.5 seconds. Twenty-five participants also completed a surprise subsequent memory test for chosen objects 24 hours following their second session; these data were not analyzed for this report.

### Behavioral analysis

On each trial, we recorded whether participants chose the option with the higher probability of correct feedback, as well as reaction time (RT) and medication state. For behavioral analysis, we divided trials into 10 learning blocks (2 blocks per scan run). We estimated a mixed effects logistic regression using *lme4* (Bates et al., 2015) with average performance and medication effect varying randomly by subject and by learning block. In *glmer* syntax for clarity:

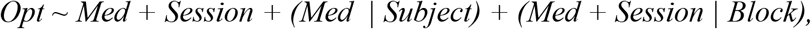

where *Med* indicates ON or OFF medication session (coded as 0.5 or −0.5) and *Session* indexes the first or second session (also coded as 0.5 or −0.5). For inference, we performed a parametric bootstrap using the varying effects estimated for learning block. To characterize task- and dopamine-related changes in reaction time, we also fit a linear mixed effects model of *log(RT)* with identical predictors and varying effects.

### Image Acquisition

Images were acquired on a 3T General Electric Signa MRI scanner using a 32-channel head coil. Functional images were acquired using a multiband pulse sequence with the following parameters: TR=850 ms, TE=25 ms, flip angle = 60°, field of view (FOV) = 192 mm. A high-resolution (1 mm isotropic) T1-weighted image was also acquired using the BRAVO pulse sequence for image co-registration. Functional runs for 5 sessions (no more than 1 run per session) were lost during acquisition due to errors in multiband image reconstruction.

### Preprocessing

Functional images were preprocessed using FSL FMRI Expert Analysis Toolbox (FEAT; (Smith et al., 2004)). Images were corrected for baseline magnetic field inhomogeneity using FUGUE. The first five images from each block were removed to account for saturation effects. The first of these five saturated images from each scan run were averaged together across blocks to form a functional template for registration. Images were high-pass filtered at *f* < 0.008 Hz, spatially smoothed with a 5 mm Gaussian kernel, grand-mean scaled, and motion corrected to the averaged template image using an affine transformation with trilinear interpolation. Due to well-characterized motion artifacts in measures of functional connectivity (Power et al. 2012; Satterthwaite et al. 2012), we used a previously validated nuisance regression strategy (Satterthwaite et al. 2013) to further preprocess functional images. Predictors in this nuisance regression included the 6 translation and rotation parameters from the motion correction registration as well as CSF, white matter, and whole-brain average time course, in addition to the square, derivative, and squared derivative of each confound. This method has been shown to outperform a number of other strategies for motion correction—including PCA- and ICA-based decomposition, global signal regression, and motion regression techniques with fewer parameters—on measures of connectivity-motion and modularity-motion correlations as well as on measures of network identifiability (Ciric et al. 2018).

Preprocessed residual images were registered to individuals’ anatomical images using linear boundary-based registration (BBR) (Greve and Fischl, 2009), and were then registered to a standard space MNI template using a nonlinear registration implemented in FNIRT. To characterize the effect of learning and dopamine on dynamic changes in functional connectivity, we used the Harvard-Oxford atlas of 110 cortical and subcortical regions (including 6 striatal regions: bilateral caudate, putamen, and nucleus accumbens) in keeping with our previous report on dynamic flexibility during reinforcement learning (Gerraty et al. 2018). After registration to 2mm MNI space, average time courses were extracted for each region in the atlas.

### Dynamic connectivity estimation

Time series were concatenated across blocks, and we used the multiplication of temporal derivatives (Shine et al., 2015) to characterize dynamic connectivity between regions. This method has been shown to be capable of detecting changes in community structure with more sensitivity than standard sliding window techniques. In this technique, dynamic coupling between each pair of regions *i* and *j* at each point in time *t* is calculated as the ratio of the product of their temporal derivatives and the product of their standard deviations. This estimate is then smoothed using a running average:

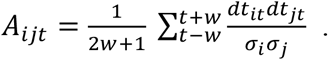

The element A_ijt_ is the dynamic functional connectivity between regions *i* and *j* at time point *t, dt_it_* is the temporal derivative, *σ_i_* is the standard deviation of the average timeseries for region *i*, and *w* is the number of time points on each side of time *t* used for smoothing. We used a window length of 13 TRs for temporal smoothing, corresponding to *w*=6 and averaging across 5.1 seconds on each side of the connectivity estimate for each time point. Using this metric, we computed a smoothed measure of network coupling for each pair of regions, leading to an N x N x T (e.g., 110 × 110 × 1477) connectivity matrix for each session. We use this matrix to represent a temporal network in which network nodes represent brain areas in the whole-brain parcellation, and network edges represent the strength of functional connectivity between pairs of regions.

### Temporal community detection

One of the most useful tools for characterizing network structure in the brain has been a set of techniques for community (or “module”) detection (Bassett et al., 2013). These techniques allow for the partitioning of the brain into internally dense, externally sparse groups of nodes based on connectivity strength. Such communities can be extracted at rest or during task performance, and provide a striking match to known cognitive systems identified and characterized through other methods. Communities in brain networks have been shown to undergo non-trivial rearrangement during reinforcement learning (Gerraty et al., 2018), as well as other cognitive and motor tasks (Bassett et al., 2011; Braun et al., 2015). To characterize the evolution of network structure in this experiment, we used a recently developed multilayer community detection algorithm (Mucha et al., 2010), which uses identity links to connect networks in neighboring time windows, in order to solve the community-matching problem and provide time-dependent labels for community assignment (Bassett et al., 2013).

Each connectivity matrix was treated as an unthresholded graph and because the graph extends in time, it represents a *temporal network*, or an ensemble of graphs ordered in time (Holme and Saramäki, 2011). Because each layer has the same number of slices, it can be represented as a *multilayer network* (Kivelä et al., 2014). Here, we partitioned regions in the multilayer network into temporal communities using a Louvain-like locally greedy algorithm for multilayer modularity maximization (Mucha et al., 2010; Jutla et al., 2011; Bassett et al., 2013). The quality function maximized in this algorithm is:

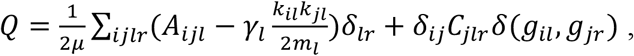

where the adjacency matrix for each layer *l* consists of pairwise connectivity components *A_iji_*. In this case, *l* is equivalent to a time point *t* in the multiplication of temporal derivatives coupling measure. The variable *γ_l_* represents the resolution parameter for layer *l*, and *C_jlr_* indexes the coupling strength between node *j* at layer *l* and node *j* at layer *r* (due to the large number of layers being decomposed into temporal communities in this study, we set both the resolution and the coupling to 1 rather than optimizing the pair of hyperparameters); *k_il_* is the coupling strength (the sum of edge weights over all connections) of node *i* in layer *l*; and *m_l_* is the average coupling strength (the sum of all edge weights over all connections for nodes *i* and *j* divided by 2). Finally, the variables *g_il_* and *g_jr_* correspond to the community labels for node *i* at layer *l* and node *j* at layer *r*, respectively, and *δ*(*g_il_, g_jr_*) is the Kronecker delta function, which equals 1 if *g_il_* = *g_jr_*, and 0 otherwise.

### Network Diagnostics

We utilized two network diagnostics to characterize dynamic connectivity changes during learning and their relationship to dopaminergic state. First, we computed the *flexibility* of each brain region, which measures the proportion of time points during which the community assignment for a node changes. This metric has been shown to relate to reinforcement learning in a previous report (Gerraty et al., 2018), as well as to motor sequence learning (Bassett et al., 2013) and executive function (Braun et al., 2015). While there are potential ambiguities in the measure related to uncertainty in node assignment, we take it to index roughly the extent to which a region is coupling with multiple networks during any given time period. We divided the community assignments into 10 blocks (2 for each task period in the scanner), and computed the flexibility for each region in each learning block.

To characterize the network connectivity of the striatum in more detail—specifically, to characterize *which* regions the striatum couples with and whether these connections are altered by dopaminergic state—we estimated the community *allegiance* for each pair of regions in each learning block. Allegiance measures the proportion of time points in a given window at which each pair of regions is assigned to the same community. The measure has been linked to motor learning (Bassett et al., 2015), as well as reinforcement learning (Gerraty et al., 2018). To disentangle domain-specific visual areas of the inferior temporal lobe, we used the Brainnetome atlas, a finer parcellation of 246 regions (Fan et al., 2016), to characterize which areas change their allegiance with the striatum during reversal learning. This atlas includes the following 12 striatal sub-regions: bilateral ventromedial and dorsolateral putamen, ventral and dorsal caudate, globus pallidus, and nucleus accumbens. For analyses of striatal allegiance, we used sub-regions overlapping with the Harvard-Oxford atlas, thus excluding the globus pallidus. Time series from each of these 246 regions underwent identical connectivity and community detection analyses to those described above.

### Flexibility Analysis

To characterize the time course of flexible striatal connectivity during learning, and the effect of choice reversal and dopamine on these dynamics, we estimated a linear mixed effects model analogous to that used to model behavior. This model predicted flexibility with estimates of average flexibility (intercept) and medication effect varying by subject, learning block, and striatal sub-region, and can be written in *lmer* syntax as:

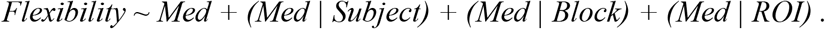

To characterize uncertainty about the time course of flexibility ON and OFF medication, we performed a parametric bootstrap using the varying intercept and medication effects estimated by learning block. To uncover regions outside of the striatum showing medication effects on network flexibility, we extended this analysis by including all Harvard-Oxford ROIs, with medication estimates varying by region and thus regularizing our estimates of the medication effect for each ROI using the distribution across brain areas. We bootstrapped intervals for these estimates in the same fashion. For both striatal and whole-brain analyses we subtracted each region’s flexibility in the first learning block and fit the above models to blocks 2-10 in order to estimate changes in flexibility from baseline during learning.

### Allegiance Analysis

To analyze changes in striatal connectivity with specific target regions, we extracted the allegiance between each striatal sub-region and every other region using the higher resolution Brainnetome atlas (Fan et al., 2016). To characterize temporal changes in striatal-cortical connectivity and the effect of dopaminergic state on these changes, we estimated the following linear mixed effects model, presented in *lmer* syntax:

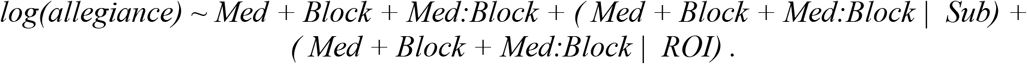

*Allegiance* is the community allegiance computed between each pair of striatal sub-regions and non-striatal target regions, *Med* is the participants’ medication state for each session, and *Block* is a factor indexing learning blocks. In this model we divided the task into 5 blocks rather than 10, corresponding in this case to scan run, due to the large number of parameters in the medication x block interaction. Effects varied randomly by participant (*Sub*) and striatal region (*ROI*). Due to the large number of regions in this more fine-grained atlas (multiplied by the number of striatal regions), we estimated this model separately for each target region and approximated a *p*-value for each term using a Wald Chi-square test, rather than allowing effects to vary randomly by target region and bootstrapping.

To evaluate the functional specificity of any time- or medication-related changes in connectivity with the striatum, we constructed a metric for correspondence to known scene- or object-processing areas for each region in the atlas, using reverse inference maps from Neurosynth (Yarkoni et al., 2011). We thresholded reverse inference maps for the terms ‘object’ and ‘place’ at Z>3.1 and binarized them to create masks. The object-scene selective metric was simply the number of voxels in a given region overlapping with the object mask minus the number of voxels overlapping with the place mask, divided by the total number of voxels in the region. This approach gave each region a weight for correspondence to object- or scene-related areas, and the weight was positive for objects and negative for scenes.

To test whether connectivity between the striatum and any domain-specific visual processing regions showed time or medication effects, we first fit the above model to regions in the top 5% of the absolute value of object-scene processing weights and corrected for multiple comparisons for each parameter using the False Discovery Rate (FDR) with q < 0.05 across these selective regions (Benjamini and Hochberg 1995). To test for the specificity of this effect to relevant scene or object processing regions, we used a more liberal threshold of p < 0.01 and calculated the average object-scene processing weight across regions passing the threshold. We then compared this average to means bootstrapped from random samples of object-scene weights from the same number of regions, taking the absolute value to capture both object and scene area correspondence. We performed both of these procedures for learning block, medication, and block x medication interaction parameters in the above model.

## Results

### Behavioral Results

Overall, participants learned to track the correct choice over the course of the experiment. We fit a mixed effects logistic regression to characterize performance and the effects of medication. Somewhat surprisingly, we did not observe a robust overall difference in performance between patients ON vs. OFF medication (parameter estimate *β* = 0.08, 95% confidence interval (CI) = [-0.03, 0.17], increase in proportion correct CI = [- 0.006, 0.04]; **Figure 2A**). There was some evidence for a small effect of dopamine in individual learning blocks, particularly before reversal (*p*(ON<OFF) < 0.05 in block 2, and *p*(ON<OFF) < 0.15 in blocks 1-5 and block 9). While Parkinson’s patients have been shown to exhibit deficits in multiple forms of feedback-based learning, and levodopa has been shown to alleviate such deficits, these effects are not always present, and some studies have found the reverse effect (Shohamy et al., 2004; Cools et al., 2007; Schonberg et al., 2010; Grogan et al., 2017; Timmer et al., 2017). We observed no effect of medication on reaction times, which decreased over the course of the task, particularly after the first block (**Figure 2B**).

**Figure 2.**
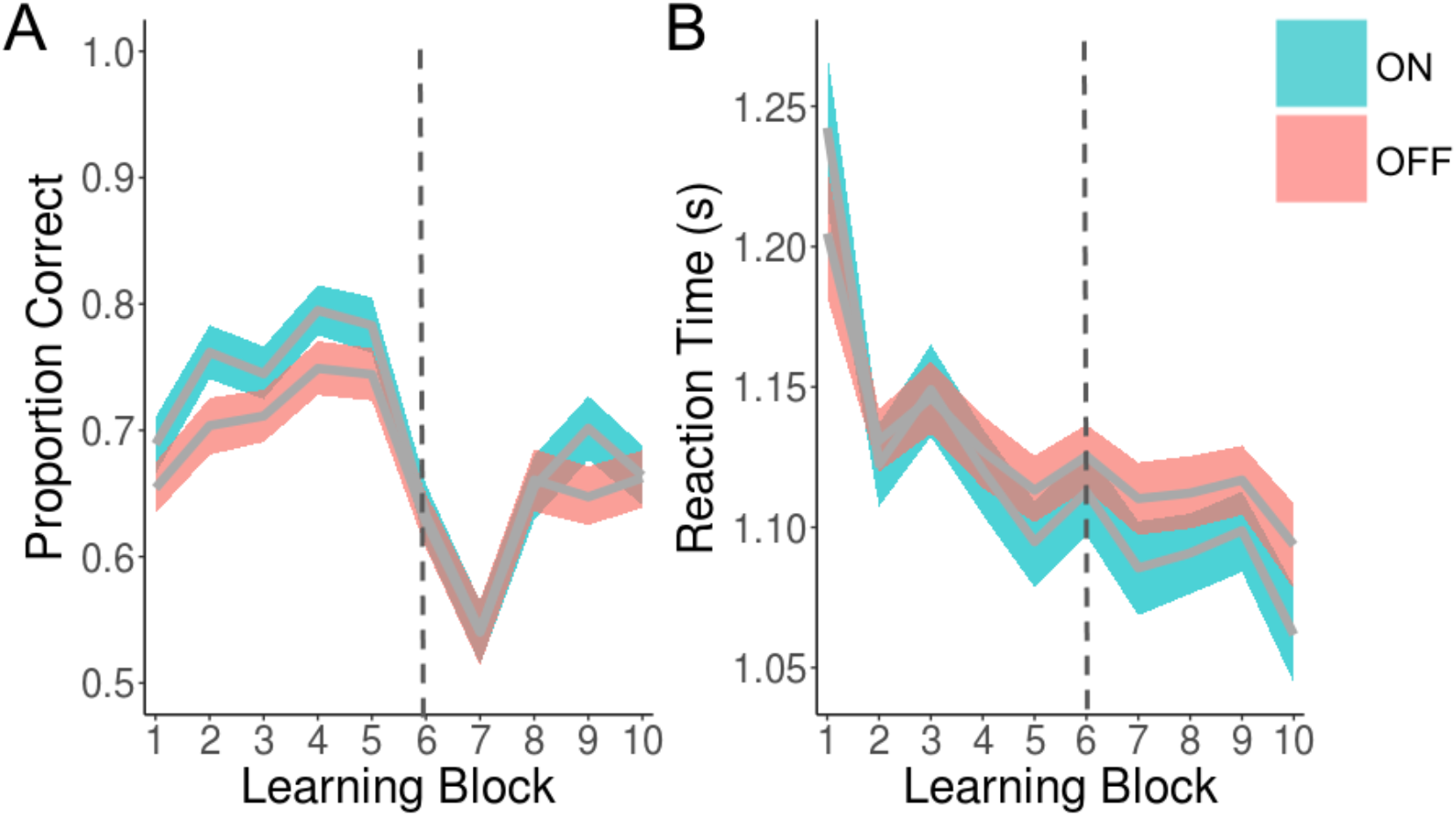
Learning in Parkinson’s patients ON and OFF of dopaminergic medication pre and post reversal. **A)** Behavioral performance, as measured by the proportion of optimal responses, is plotted against trials binned into 10 learning blocks. The grey lines indicate the average proportion of correct choices and the ribbons indicate standard errors, estimated by bootstrapping a generalized logistic mixed effects model of optimal choice. **B)** Reaction time (RT, seconds) is plotted against trials binned into the same 10 learning blocks. Grey lines indicate the geometric mean RT and the ribbons indicate standard errors, estimated by bootstrapping a linear mixed effects model of log RT. Colors show medication state (ON levodopa = blue, OFF levodopa = red). The point at which the reversal occurred is indicated by dotted lines in both panels.

### Learning and dopamine both modulate dynamic flexibility in the striatum

Previous studies of flexibility have been limited in interpretation by the fact that they have relied on correlations with behavioral measures. We designed this task with a reversal in order to provide an experimental manipulation of this network metric. We extracted time series from regions distributed across the brain and computed the dynamic connectivity between all pairs of regions at each time point during the task. We summarized dynamic connectivity in the form of temporal multilayer networks and extracted the reconfiguration of functional modules using a multilayer community detection technique. After splitting the task into 10 learning blocks per session (2 per scanner run), we calculated the *network flexibility* in each block to capture the dynamic connectivity of regions during different periods of the task. We were particularly interested in flexibility in striatal connectivity, as we have previously shown this network metric to relate to reinforcement learning (Gerraty et al. 2018).

To test the effects of dopamine on dynamic striatal flexibility, we fit a linear mixed effects model with average flexibility and medication state varying by subject and by striatal sub-region. In both the ON and OFF medication conditions, network flexibility in the striatum decreases as a result of reversal before recovering, mirroring changes in learning performance (**Figure 3A**). Overall flexibility was higher in the OFF medication condition than in the ON medication condition (regression *β* = −0.004, CI = [-0.006, - 0.002], *p*(ON<OFF) < 0.01), potentially due to medication-related differences in motion or in overall differences in dynamic brain connectivity. Because we were most interested in *changes* in network flexibility over the course of learning, we subtracted the flexibility measured in the first learning bin from the flexibility measured in all other bins. As can be seen in **Figure 3B**, this baseline-subtracted flexibility measure was higher in patients ON dopamine medication than OFF dopamine medication (regression *β* = 0.01, CI = [0.009, 0.02], *p*(ON<OFF) < 0.01), indicating greater modulation of striatal-network coupling in the ON dopamine condition. This effect was similar across sub-regions of the striatum (**Figure 3C**).

**Figure 3.**
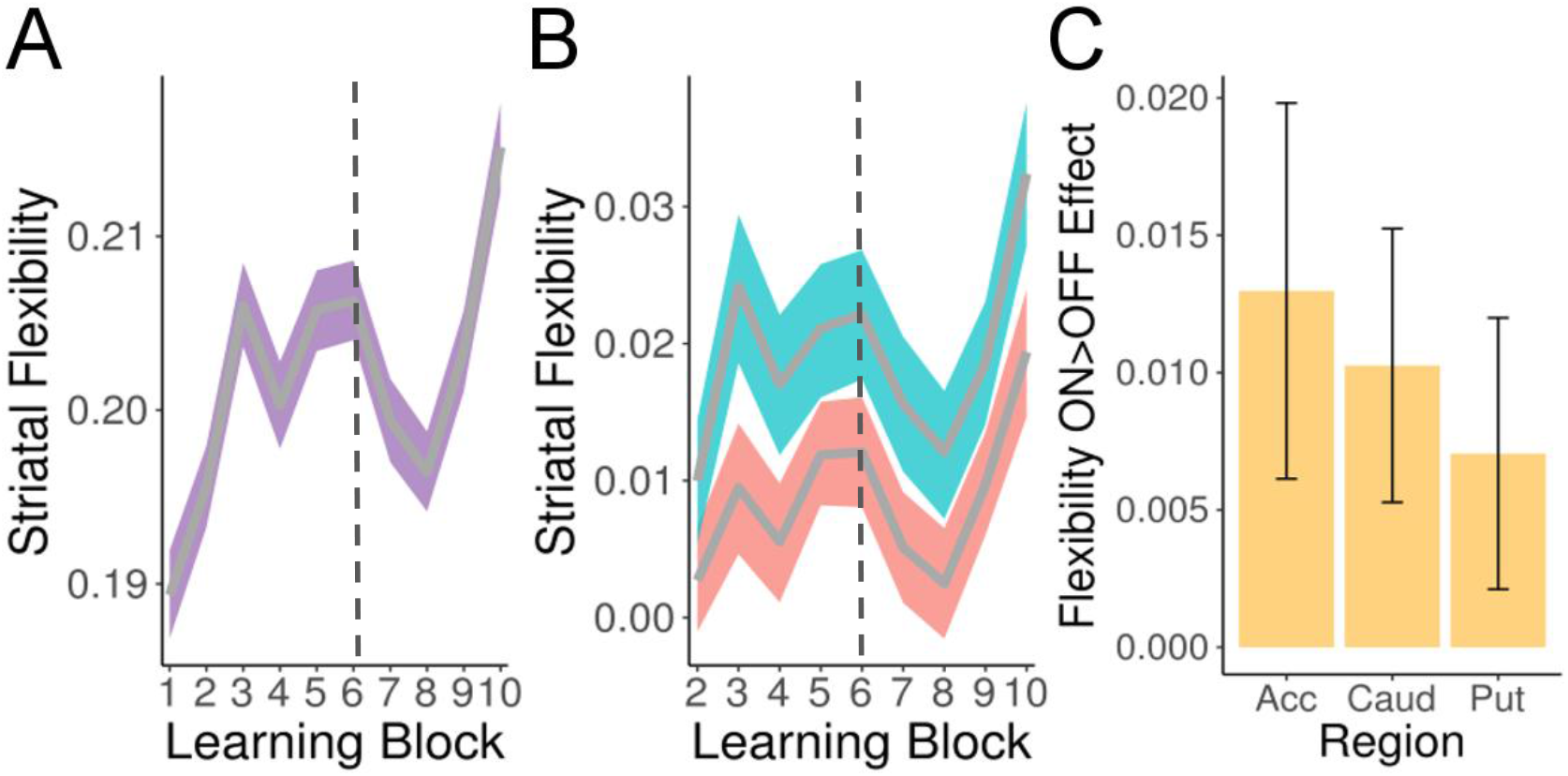
Learning-related changes in dynamic striatal connectivity are modulated by dopamine. **A)** Reversal in contingencies is reflected in changes in network flexibility in the striatum. Striatal flexibility, as measured by the proportion of network changes in a time window, is plotted against learning block. The reversal of outcome contingencies provided an experimental manipulation of flexibility, which decreased following reversal before recovering, mirroring behavioral performance (**Figure 2**). **B)** Dopaminergic state modulates changes in striatal flexibility. Plot shows baseline-subtracted network flexibility in the striatum over learning blocks. Grey lines indicate the mean flexibility and the ribbons indicate standard errors, estimated by bootstrapping a linear mixed effects model of flexibility varying by subject ROI, learning block, and striatal sub-region. The dotted lines in panels A and B illustrate the reversal of outcome contingencies in the 6th learning block. C. The effect of medication was similar across sub-regions of the striatum. Across all panels, colors show medication state (ON levodopa = blue, OFF levodopa = red, average across states = purple). Bars show parameter estimates for separate models fit to each ROI; error bars indicate standard errors. Acc = Nucleus Accumbens, Caud = Caudate, and Put = Putamen.

### Dopamine has widespread effects on network flexibility

Dopamine has widespread cortical and subcortical targets, and we have previously shown that network flexibility in regions of parietal and prefrontal cortex also relates to reinforcement learning. Thus, we were interested in whether other regions exhibited greater flexibility in network coupling ON versus OFF dopamine medication. To examine this question, we estimated a linear mixed effects model with medication effects in network flexibility varying randomly by subject, learning block, and brain region. Region-level differences surpassing p<0.01 included the nucleus accumbens, hippocampus, ventromedial and dorsolateral prefrontal cortex, as well as supplementary motor and parietal cortex (**Figure 4**). All regions passing this threshold exhibited greater flexibility ON dopamine medication relative to OFF dopamine medication. Given its anatomical position as a major target of midbrain dopamine, and its established role in value-based learning, we also extracted the time course of flexibility from the ventromedial prefrontal cortex (vmPFC) region showing a medication effect. As seen in **Figure 4B**, flexibility in this region was also affected by reversal, further supporting a link between dopamine, learning, and changes in network connectivity.

**Figure 4.**
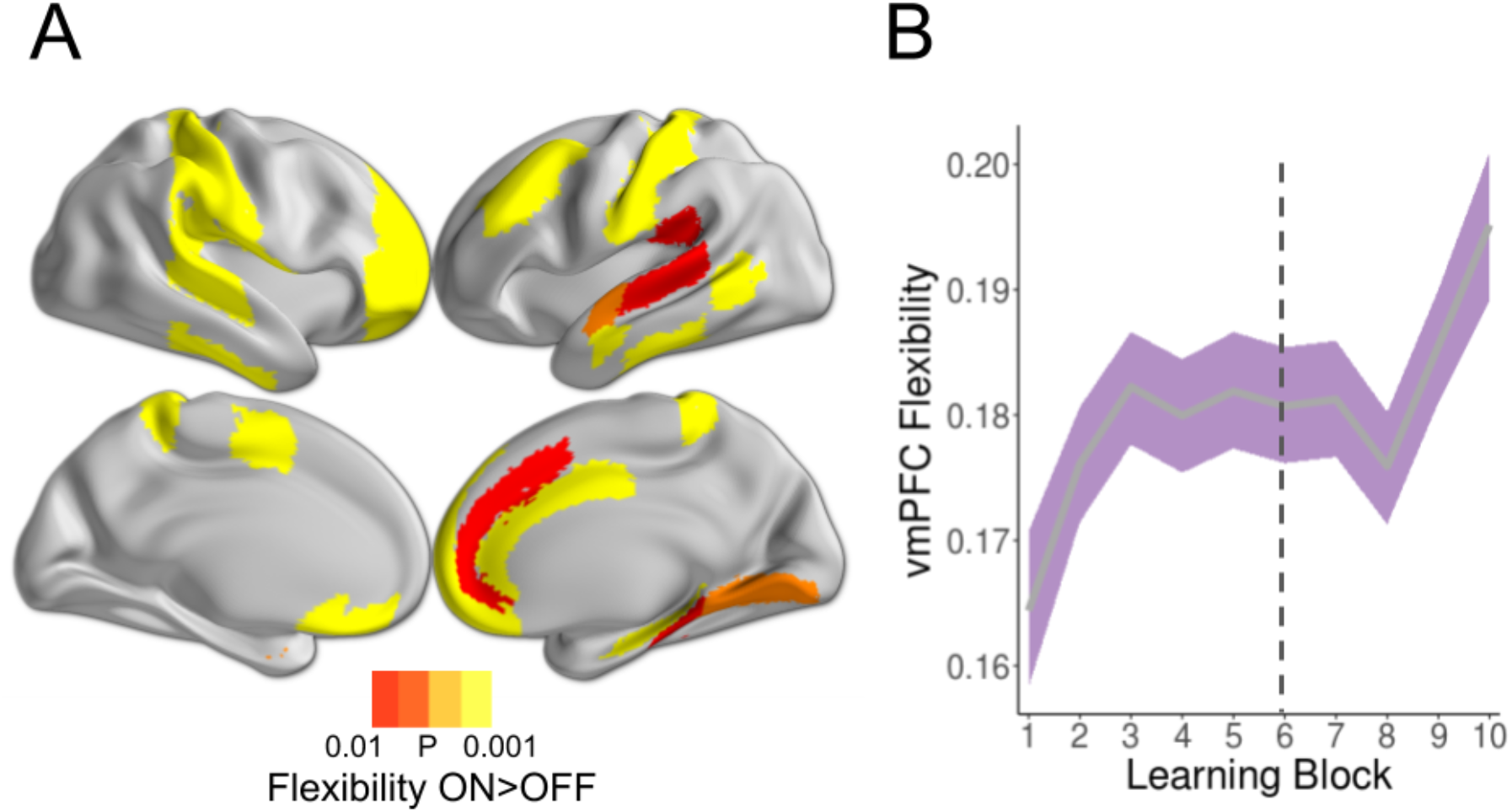
Dopamine increases dynamic connectivity across multiple brain areas. **A)** The hippocampus, vmPFC, and regions across limbic, parietal, and prefrontal cortex exhibit greater learning-related changes in dynamic connectivity ON dopamine medication relative to OFF dopamine medication. The color overlay displays results from a linear mixed effects model of baseline-subtracted flexibility, with medication effect varying by subject, learning block, and ROI. Medication effects were parametrically bootstrapped from the varying ROI estimates, and thresholded at *p*(ON<OFF) < 0.01. Lighter color indicates lower *p*(ON<OFF). B) An ROI analysis of ventromedial prefrontal cortex (vmPFC) reveals that flexibility in this region is also modulated by the reversal task. The grey line indicates the mean flexibility and the ribbons indicate standard errors, estimated by bootstrapping a linear mixed effects model of vmPFC flexibility.

### Dopamine modulates dynamic coupling between the striatum and task-specific visual areas

Having shown that dynamic connectivity in the striatum changes over the course of learning and is modulated by dopamine, we sought to determine *which* regions vary in their striatal coupling. The task was designed to capture the effect of dopamine on dynamic coupling between the striatum and areas of visual cortex. In particular, we hypothesized that the striatum would exhibit greater changes in connectivity with domain-specific areas that contribute to the processing of scene or object stimuli, and that these changes would be affected by dopaminergic state.

To capture these specific functional regions of cortex, we used a more fine-grained atlas of 246 regions (Fan et al., 2016). We constructed a measure of correspondence between each region in this atlas and areas contributing to the processing of scene or object information using Neurosynth (Yarkoni et al., 2011); see **Methods**. We calculated the *community allegiance* between the striatum and each of these regions. To test whether category-specific visual regions exhibited dynamic changes in connectivity with the striatum during the task, and whether any such changes were affected by dopamine, we estimated a linear mixed effects model of changes in striatal allegiance with regions showing the strongest (top 5%) correspondence to scene- or object-specific areas. In this model, we included learning block, medication state, and the interaction between the two.

No regions showed an overall effect of medication on average striatal allegiance. A region in the right fusiform gyrus with a higher correspondence to object areas exhibited a significant effect of learning block (*p*<0.05 FDR corrected; **Figure 5A**, top), as well as a significant dopaminergic modulation of these temporal changes (*p*<0.05 FDR corrected; **Figure 5B**).

**Figure 5.**
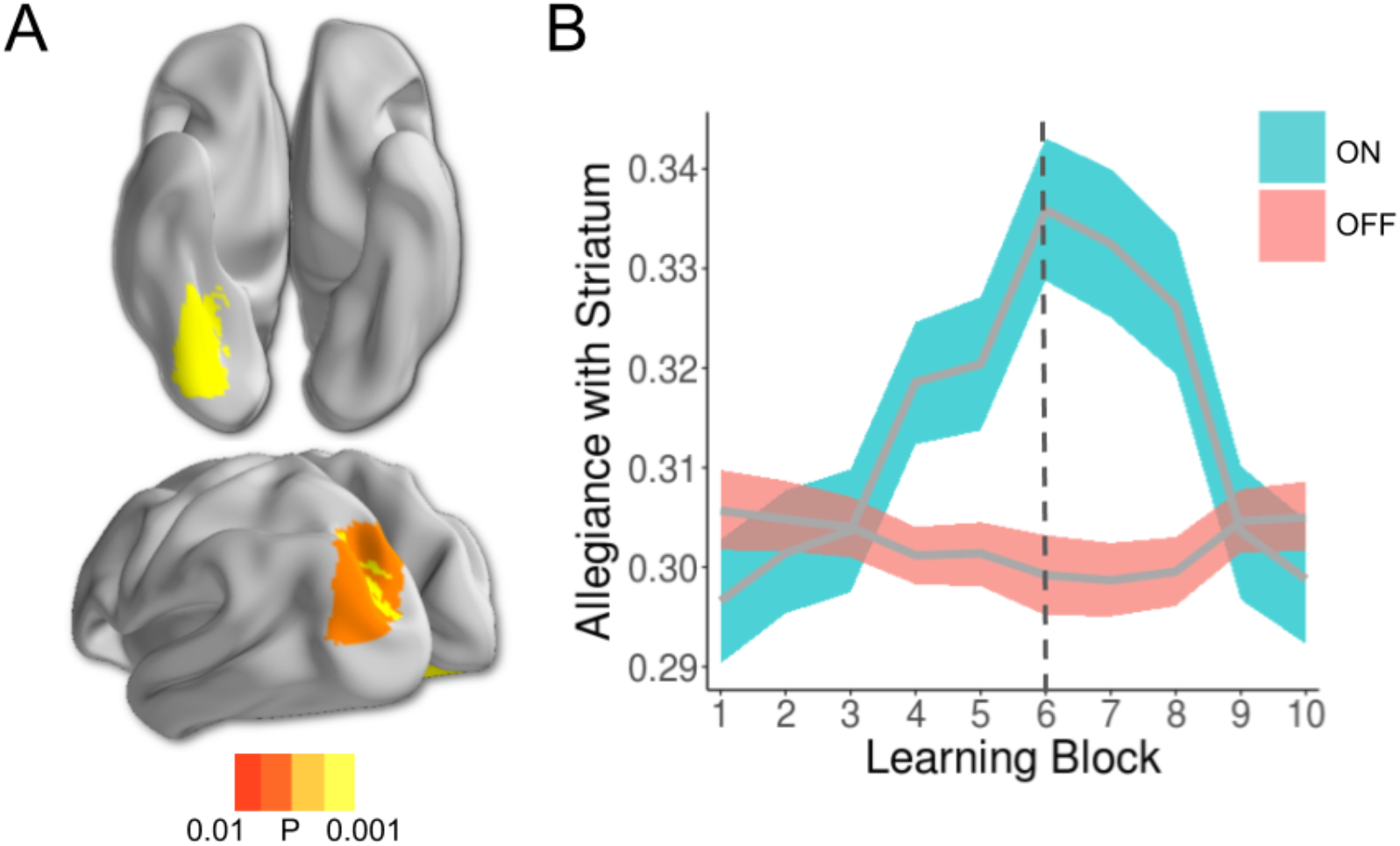
Dopamine modulates dynamic connectivity between the striatum and task-relevant visual regions. **A)** Regions exhibiting changes in module allegiance with the striatum over the course of the task. Shown at a threshold of *p* < 0.01, these regions include the fusiform gyrus (*top*), inferior parietal lobe (*bottom*), and lateral superior occipital gyrus (*bottom*). Based on Neurosynth maps, these regions showed a greater correspondence to object-processing areas than would be expected by chance alone. B) Interaction between medication state and changes in allegiance in the fusiform gyrus region shown in panel A (*top*). This area of inferior temporal cortex showed significant temporal changes in striatal allegiance, as well as a significant dopaminergic effect on these changes (both FDR *p*<0.05). Solid lines show mean allegiance with the striatum, and ribbons show standard error. Colors indicate medication group (blue=ON, red=OFF).

This result demonstrates that dynamic connectivity between the striatum and a category-specific visual region is altered by dopamine, but the analysis was limited to visual areas. To test the specificity of these effects, we used a lower threshold (*p*<0.01, uncorrected) across all brain regions. Two more regions in occipital and inferior parietal lobe (IPL) passed this more liberal threshold for an effect of time on striatal allegiance (**Figure 5A**, bottom), and the same IPL region passed this threshold for a significant dopamine-by-time interaction. These regions showed a significantly higher level of object correspondence than would be expected due to chance alone (*p*=0.014 for the effect of learning block; *p* < 0.005 for the block-by-medication interaction; see **Methods** for description of bootstrapping procedure). Collectively, these results provide further evidence of a dopaminergic effect on the dynamics of striatal connectivity with task-relevant regions during learning.

## Discussion

There is growing evidence that dynamic changes in cortico-striatal connectivity play an important role in value-based learning (Mattar et al., 2016; Gerraty et al., 2018). Here, we show that changes in striatal connectivity are modulated by the neurotransmitter dopamine, which has been theorized to facilitate learning by updating associations via changes in striatal-cortical connections (White, 1989b; Glimcher, 2011). In addition, by experimentally manipulating learning with a reversal of contingencies and showing corresponding changes in dynamic cortico-striatal connectivity, our results lend causal support to previously demonstrated associations between network flexibility in the striatum and learning. Our finding that dopamine-induced changes in cortico-striatal coupling are most pronounced in task-relevant domain-specific sensory regions provides further empirical support for this theory.

Dopamine is known to play an essential role in reinforcement learning. Evidence of its importance has come from experimental dopaminergic manipulation in non-human animals (Beninger, 1983), studies in healthy controls (Pizzagalli et al., 2008), and studies in patients with Parkinson’s disease (Knowlton et al., 1996; Rutledge et al., 2009; Shiner et al., 2012; Schmidt et al., 2014; Sharp et al., 2015). Reinforcement learning, in turn, involves coordinated activation across widespread brain circuits, with recent work linking dynamic connectivity to reward or value learning. Given dopamine’s widespread cortical innervation and its well-established role in learning, it is a likely candidate for involvement in coordinating large-scale network dynamics. The findings reported here link this essential neurostransmitter to learning-related changes in dynamic network structure.

Our findings are particularly interesting in light of the reward prediction error theory of midbrain dopamine function. It has been shown that dopaminergic neurons in the midbrain fire in a manner consistent with the temporal difference errors postulated in reinforcement learning algorithms (Schultz et al., 1997; Bayer and Glimcher, 2005). These findings have been widely influential in neuroscientific theories of learning, providing a parsimonious account of dopamine’s role in the process. And while it is not precisely known how this dopaminergic signal affects learned associations, it has been suggested that prediction errors update associations through changes in synaptic weights at widespread cortical targets of dopamine that are active following a given state and action (Glimcher, 2011). This theory is intuitive, consistent with known anatomy, and has proved difficult to test empirically. So, most studies to date have focused on the activity of single neurons or brain regions.

Network neuroscience has provided a wealth of experimental evidence that the characteristics of distributed brain circuits are relevant to cognition (see (Medaglia et al., 2015; Petersen and Sporns, 2015; Bassett and Sporns, 2017; Bassett et al., 2018) for recent reviews). However, measures that can be used to characterize brain networks have been hampered by an inability to describe dynamic changes in these circuits. Our report takes advantage of recent developments in network science that afford precisely this ability (Bassett et al., 2013; Khambhati et al., 2017; Sizemore and Bassett, 2017): we show that cortico-striatal circuits change as a result of reinforced associations, and that these changes are modulated by dopamine.

Our behavioral findings are more ambiguous, which is consistent with previous studies of reversal learning in Parkinson’s patients. While patients are known to exhibit subtle deficits in learning from feedback, and while some research has shown improved performance on dopaminergic medication, other studies have failed to find medication-related improvement on reversal tasks (Cools et al., 2007; Shohamy et al., 2009). We did not observe robust differences in performance or reaction time, although patients did perform slightly better on medication. This could be due to counterbalancing between medication state and session (such that half of the off medication sessions were measured after participants had already performed the task on medication) or to the task contingencies being too easy for patients to learn. It also raises the possibility that distinct cognitive strategies and neural processes could lead to behavioral results that appear similar on the surface. But with a single reversal and counterbalanced design, caution should be exercised in interpreting the lack of robust behavioral differences.

This report has a number of limitations. First, the multilayer community detection algorithm we used provides deterministic community labels, and is unable to assess the possibility of regions probabilistically coupling with multiple communities at a given time point. Addressing this limitation may be important for providing a fuller account of dynamic changes in brain networks during learning and other cognitive processes. The further development of probabilistic and generative models, which poses serious statistical and computational challenges, will be essential in this regard (Durante et al., 2016; Palla et al., 2016; Betzel and Bassett, 2017). Second, while our results are consistent with the prediction error account of dopamine function, we did not have the temporal resolution to characterize dynamic network changes at the level of individual trials, which would be necessary to link network topology directly to prediction error updating. Indeed, the use of fMRI, which affords a crucial level of spatial specificity, limits the temporal resolution with which we can describe brain network dynamics. Linking individual trial level updating with network flexibility will be an important area for future studies. In addition, to better characterize network changes underlying learning, it may be useful to measure the effect of individual prediction errors on network structure, and network studies using ECoG or combining fMRI and EEG may prove useful in this regard.

A further limitation to this study is the use of a patient group with no control participants. We chose to focus on a within-subject pharmacological manipulation so as to limit the number of terms in the statistical interactions necessary to characterize dopaminergic effects, thereby limiting the uncertainty in our inferences. It is possible that these results are specific to patients with Parkinson’s disease; however, this concern may be mitigated by the fact that patients learned the correct choice and updated their decisions following reversal. Nevertheless, further research could usefully evaluate similar hypotheses in neurologically intact humans.

While network science has produced a growing number of results linking large-scale brain circuits to cognitive processes such as learning, it is sometimes unclear what the theoretical implications of these results are. In this study we sought to characterize, based on convergence between previous studies of dynamic network changes during learning and computational theories of reinforcement, the effect of dopamine on learning-related changes in striatal-cortical coupling. In addition to providing experimental support linking dynamic connectivity to feedback-based learning and a neurotransmitter system known to underlie this process, this study illustrates the potential utility of integrating network neuroscience with more established theoretic frameworks for understanding cognition.

## Acknowledgements

The authors thank Christina Galese for patient recruitment, testing, and imaging acquisition. Thanks to Camilla van Geen for helpful comments on previous drafts. This work was supported by a New York State Psychiatric Institute MRI Pilot Award. R.T.G. acknowledges support from the National Institute of Mental Health (F31 MH109247-01A1). M.E.S. acknowledges support from the Canadian Institutes of Health Research Fellowship and the Paul Janssen Fellowship in Translational Neuroscience. D.S.B. acknowledges support from the John D. and Catherine T. MacArthur Foundation, the Alfred P. Sloan Foundation, the ISI Foundation, the Paul Allen Foundation, the Army Research Laboratory (W911NF-10-2-0022), the Army Research Office (W911NF-14-1-0679, W911NF-16-1-0474), the National Institute of Mental Health (2-R01-DC-009209-11, R01 - MH112847, R01-MH107235, R21-M MH-106799), the National Institute of Child Health and Human Development (1R01HD086888-01), National Institute of Neurological Disorders and Stroke (R01 NS099348), and the National Science Foundation (BCS-1441502, BCS-1430087, NSF PHY-1554488 and BCS-1631550). D.S. acknowledges support from the National Institute on Drug Abuse (R01DA038891).

## References

Bassett DS, Sporns O (2017) Network neuroscience. Nature neuroscience 20:353.

Bassett DS, Zurn P, Gold JI (2018) On the nature and use of models in network neuroscience. Nature Reviews Neuroscience: 1.

Bassett DS, Yang M, Wymbs NF, Grafton ST (2015) Learning-induced autonomy of sensorimotor systems. Nature Neuroscience.

Bassett DS, Porter MA, Wymbs NF, Grafton ST, Carlson JM (2013) Robust detection of dynamic community structure in networks. Chaos 23:013142.

Bassett DS, Wymbs NF, Porter MA, Mucha PJ, Carlson JM, Grafton ST (2011) Dynamic reconfiguration of human brain networks during learning. Proceedings of the National Academy of Sciences 108:7641–7646.

Bates D, Mächler M, Bolker B, Walker S (2015) Fitting Linear Mixed-Effects Models Usinglme4. Journal of Statistical Software 67.

Bayer HM, Glimcher PW (2005) Midbrain dopamine neurons encode a quantitative reward prediction error signal. Neuron 47:129–141.

Beninger RJ (1983) The role of dopamine in locomotor activity and learning. Brain Research Reviews 6:173–196.

Betzel RF, Bassett DS (2017) Generative models for network neuroscience: prospects and promise. Journal of The Royal Society Interface 14:20170623.

Bogacz R, Gurney K (2007) The basal ganglia and cortex implement optimal decision making between alternative actions. Neural computation 19:442–477.

Braun U, Schäfer A, Walter H, Erk S, Romanczuk-Seiferth N, Haddad L, Schweiger JI, Grimm O, Heinz A, Tost H (2015) Dynamic reconfiguration of frontal brain networks during executive cognition in humans. Proceedings of the National Academy of Sciences 112:11678–11683.

Cools R, Lewis SJ, Clark L, Barker RA, Robbins TW (2007) L-DOPA disrupts activity in the nucleus accumbens during reversal learning in Parkinson’s disease. Neuropsychopharmacology 32:180.

Daw ND, O’Doherty JP, Dayan P, Seymour B, Dolan RJ (2006) Cortical substrates for exploratory decisions in humans. Nature 441:876–879.

Ding L (2015) Distinct dynamics of ramping activity in the frontal cortex and caudate nucleus in monkeys. Journal of neurophysiology 114:1850–1861.

Durante D, Mukherjee N, Steorts RC (2016) Bayesian Learning of Dynamic Multilayer Networks. arXiv preprint arXiv:160802209.

Epstein R, Kanwisher N (1998) A cortical representation of the local visual environment. Nature 392:598.

Epstein R, Harris A, Stanley D, Kanwisher N (1999) The parahippocampal place area: Recognition, navigation, or encoding? Neuron 23:115–125.

Fan L, Li H, Zhuo J, Zhang Y, Wang J, Chen L, Yang Z, Chu C, Xie S, Laird AR (2016) The human brainnetome atlas: a new brain atlas based on connectional architecture. Cerebral cortex 26:3508–3526.

Foerde K, Braun EK, Shohamy D (2012) A trade-off between feedback-based learning and episodic memory for feedback events: evidence from Parkinson’s disease. Neurodegenerative Diseases 11:93–101.

Frank MJ, Seeberger LC, O’reilly RC (2004) By carrot or by stick: cognitive reinforcement learning in parkinsonism. Science 306:1940–1943.

Gerraty RT, Davidow JY, Foerde K, Galvan A, Bassett DS, Shohamy D (2018) Dynamic flexibility in striatal-cortical circuits supports reinforcement learning. Journal of Neuroscience 38:2442–2453.

Glimcher PW (2011) Understanding dopamine and reinforcement learning: the dopamine reward prediction error hypothesis. Proceedings of the National Academy of Sciences 108:15647–15654.

Greve DN, Fischl B (2009) Accurate and robust brain image alignment using boundary-based registration. Neuroimage 48:63–72.

Grogan JP, Tsivos D, Smith L, Knight BE, Bogacz R, Whone A, Coulthard EJ (2017) Effects of dopamine on reinforcement learning and consolidation in Parkinson’s disease. eLife 6.

Hikosaka O, Kim HF, Yasuda M, Yamamoto S (2014) Basal ganglia circuits for reward value-guided behavior. Annual review of neuroscience 37:289.

Holme P, Saramäki J (2011) Temporal Networks. Physics Reports 519:97–125.

Jutla IS, Jeub LG, Mucha PJ (2011) A generalized Louvain method for community detection implemented in MATLAB. URL http://netwiki amath uncedu/GenLouvain.

Kao K, Powell D (1986) Lesions of substantia nigra retard Pavlovian somatomotor learning but do not affect autonomic conditioning. Neuroscience letters 64:1–6.

Kemp JM, Powell T (1971) The connexions of the striatum and globus pallidus: synthesis and speculation. Philosophical Transactions of the Royal Society of London B: Biological Sciences 262:441–457.

Khambhati AN, Sizemore AE, Betzel RF, Bassett DS (2017) Modeling and interpreting mesoscale network dynamics. Neuroimage.

Kivelä M, Arenas A, Barthelemy M, Gleeson JP, Moreno Y, Porter MA (2014) Multilayer networks. Journal of Complex Networks 2:203–271.

Knowlton BJ, Mangels JA, Squire LR (1996) A neostriatal habit learning system in humans. Science 273:1399–1402.

Mattar MG, Thompson-Schill SL, Bassett DS (2016) The network architecture of value learning. Network Neuroscience:1–27.

Medaglia JD, Lynall M-E, Bassett DS (2015) Cognitive Network Neuroscience. Journal of cognitive neuroscience.

Mucha PJ, Richardson T, Macon K, Porter MA, Onnela J-P (2010) Community structure in time-dependent, multiscale, and multiplex networks. Science 328:876–878.

Pagnoni G, Zink CF, Montague PR, Berns GS (2002) Activity in human ventral striatum locked to errors of reward prediction. Nature neuroscience 5:97.

Palla K, Caron F, Teh YW (2016) Bayesian nonparametrics for Sparse Dynamic Networks. arXiv preprint arXiv:160701624.

Palminteri S, Lebreton M, Worbe Y, Grabli D, Hartmann A, Pessiglione M (2009) Pharmacological modulation of subliminal learning in Parkinson’s and Tourette’s syndromes. Proceedings of the National Academy of Sciences 106:19179–19184.

Petersen SE, Sporns O (2015) Brain networks and cognitive architectures. Neuron 88:207–219.

Pizzagalli DA, Evins AE, Schetter EC, Frank MJ, Pajtas PE, Santesso DL, Culhane M (2008) Single dose of a dopamine agonist impairs reinforcement learning in humans: behavioral evidence from a laboratory-based measure of reward responsiveness. Psychopharmacology 196:221–232.

Reddy L, Kanwisher N (2006) Coding of visual objects in the ventral stream. Current opinion in neurobiology 16:408–414.

Robbins TW, Brown VJ (1990) The role of the striatum in the mental chronometry of action: a theoretical review. Reviews in the Neurosciences 2:181–214.

Rutledge RB, Lazzaro SC, Lau B, Myers CE, Gluck MA, Glimcher PW (2009) Dopaminergic drugs modulate learning rates and perseveration in Parkinson’s patients in a dynamic foraging task. Journal of Neuroscience 29:15104–15114.

Schmidt L, Braun EK, Wager TD, Shohamy D (2014) Mind matters: placebo enhances reward learning in Parkinson’s disease. Nature neuroscience 17:1793.

Schonberg T, O’Doherty JP, Joel D, Inzelberg R, Segev Y, Daw ND (2010) Selective impairment of prediction error signaling in human dorsolateral but not ventral striatum in Parkinson’s disease patients: evidence from a model-based fMRI study. Neuroimage 49:772–781.

Schultz W, Dayan P, Montague PR (1997) A neural substrate of prediction and reward. Science 275:1593–1599.

Sharp ME, Foerde K, Daw ND, Shohamy D (2015) Dopamine selectively remediates ‘model-based’reward learning: a computational approach. Brain 139:355–364.

Shine JM, Koyejo O, Bell PT, Gorgolewski KJ, Gilat M, Poldrack RA (2015) Estimation of dynamic functional connectivity using Multiplication of Temporal Derivatives. NeuroImage 122:399–407.

Shine JM, Bissett PG, Bell PT, Koyejo O, Balsters JH, Gorgolewski KJ, Moodie CA, Poldrack RA (2016) The Dynamics of Functional Brain Networks: Integrated Network States during Cognitive Task Performance. Neuron 92:544–554.

Shiner T, Seymour B, Wunderlich K, Hill C, Bhatia KP, Dayan P, Dolan RJ (2012) Dopamine and performance in a reinforcement learning task: evidence from Parkinson’s disease. Brain:aws083.

Shohamy D, Myers C, Onlaor S, Gluck M (2004) Role of the basal ganglia in category learning: how do patients with Parkinson’s disease learn? Behavioral neuroscience 118:676.

Shohamy D, Myers CE, Hopkins RO, Sage J, Gluck MA (2009) Distinct hippocampal and basal ganglia contributions to probabilistic learning and reversal. Journal of Cognitive Neuroscience 21:1820–1832.

Sizemore AE, Bassett DS (2017) Dynamic graph metrics: Tutorial, toolbox, and tale. NeuroImage.

Skinner BF (1948) ‘Superstition’in the pigeon. Journal of experimental psychology 38:168.

Smith SM, Jenkinson M, Woolrich MW, Beckmann CF, Behrens TE, Johansen-Berg H, Bannister PR, De Luca M, Drobnjak I, Flitney DE (2004) Advances in functional and structural MR image analysis and implementation as FSL. Neuroimage 23:S208–S219.

Steinberg EE, Keiflin R, Boivin JR, Witten IB, Deisseroth K, Janak PH (2013) A causal link between prediction errors, dopamine neurons and learning. Nature neuroscience 16:966.

Sutton RS, Barto AG (1998) Reinforcement learning: An introduction: MIT press.

Thorndike E (1898) Some experiments on animal intelligence. Science:818–824.

Timmer MH, Sescousse G, van der Schaaf ME, Esselink RA, Cools R (2017) Reward learning deficits in Parkinson’s disease depend on depression. Psychological medicine 47:2302–2311.

Vo K, Rutledge RB, Chatterjee A, Kable JW (2014) Dorsal striatum is necessary for stimulus-value but not action-value learning in humans. Brain 137:3129–3135.

Westerink B, Kwint H (1996) The pharmacology of mesolimbic dopamine neurons: a dual-probe microdialysis study in the ventral tegmental area and nucleus accumbens of the rat brain. Journal of Neuroscience 16:2605–2611.

White NM (1989a) A functional hypothesis concerning the striatal matrix and patches: Mediation of S □ R memory and reward. Life sciences 45:1943–1957.

White NM (1989b) Reward or reinforcement: what’s the difference? Neuroscience & Biobehavioral Reviews 13:181–186.

Yarkoni T, Poldrack RA, Nichols TE, Van Essen DC, Wager TD (2011) Large-scale automated synthesis of human functional neuroimaging data. Nature methods 8:665.

Yin HH, Knowlton BJ (2006) The role of the basal ganglia in habit formation. Nature Reviews Neuroscience 7:464.

Yin HH, Knowlton BJ, Balleine BW (2006) Inactivation of dorsolateral striatum enhances sensitivity to changes in the action–outcome contingency in instrumental conditioning. Behavioural brain research 166:189–196.

